# Liver-specific Inflammatory Signatures Predict Clinically Significant Liver Damage

**DOI:** 10.1101/2023.02.28.530386

**Authors:** Conan Chua, Deeqa Mahamed, Shirin Nkongolo, Aman Mehrotra, David K.H. Wong, Raymond T. Chung, Jordan J. Feld, Harry L.A. Janssen, Adam J. Gehring

## Abstract

**Background and Aims:** Inflammation drives progression of chronic liver disease. However, the triggers of inflammation remain undefined during chronic hepatitis B (CHB) because hepatic flares are spontaneous and difficult to capture. We used nucleoside analogue (NA) withdrawal to investigate early inflammatory events because liver damage after stopping therapy occurs in a predictable time frame. 11 CHB patients underwent 192 weeks of NA therapy before a protocol defined stop. Liver fine-needle aspirates (FNAs) were collected at baseline and 4-weeks post-withdrawal and analyzed using flow cytometry and single-cell RNA sequencing (scRNA-seq). Intrahepatic mononuclear cells (IHMCs) from uninfected livers were used to validate transcriptomic findings. At 4 weeks post NA-withdrawal, HBV DNA rebounded but alanine aminotransferase (ALT) levels remained normal, 7/11 patients developed ALT elevations (>2xULN) at later timepoints. There were no changes in cell frequencies between baseline and viral rebound. ScRNA-seq revealed upregulation of IFN stimulated genes (ISGs) and pro-inflammatory cytokine *MIF* upon viral rebound. In vitro experiments confirmed the type I IFN-dependent ISG profile whereas *MIF* was induced primarily by IL-12. MIF exposure further amplified inflammatory cytokine production by myeloid cells. Our data show that innate immune activation is detectable in the liver before clinically-significant liver damage is detectable in the serum.

## INTRODUCTION

Chronic Hepatitis B virus (HBV) infection affects 292 million individuals globally and approximately 800,000 deaths occur annually. The majority of HBV-associated deaths are the result of progressive liver damage leading to cirrhosis and liver cancer. HBV itself is non-cytopathic and liver damage is primarily immune-mediated. Liver damage is characterized by intrahepatic immune infiltration^1–6^, resulting from elevated chemokine and cytokine levels^7–9^. The inflammatory environment leads to activation of cytotoxic cells that mediate hepatocyte killing through an antigen-independent mechansism^7,9–11^. Hepatocyte killing then leads to the elevation of alanine amino transferase (ALT) levels, a marker of hepatocyte cell death, in the serum. The challenge to define the cascade of events leading to liver damage is that hepatic flares occur unpredictably. They are typically identified on routine patient clinic visits every 3 – 6 months, at which time, inflammation associated with active liver damage, and the initial triggers responsible for the inflammatory cascade, are indistinguishable from the inflammatory liver environment^11^. Thus, the initial triggers of the inflammatory cascade remain undefined in the human liver. Characterizing the initial drivers of liver inflammation could help to understand how early immune events evolve into potentially life-threatening liver damage.

Currently, liver damage in CHB patients is managed with nucleos(t)ide analogue (NA) therapy. NA therapy suppresses viral replication by inhibiting reverse transcription. NA therapy reduces viral load in the serum and reduces liver damage but does not eliminate the virus from the liver, meaning that patients require long-term and potentially life-long treatment. In recent years, NA discontinuation has been used as a strategy to limit the duration of NA therapy. The hypothesis being that rebound in viral replication would “shock” the immune system and lead to control of HBV replication without the need for further treatment. However, NA withdrawal is accompanied with high rates of virological and clinical relapses, even after years of treatment^12,13^. The typical course of events following NA cessation include viral rebound (defined as HBV DNA >2,000 IU/mL) within 4 weeks of withdrawal in almost all patients, followed by ALT elevation 8-18 weeks post-withdrawal that occurs in approximately 40% of HBeAg-negative patients^12,14,15^. The predictable timeline of viral rebound and liver damage provides an opportunity to investigate intrahepatic immune activation before the onset of detectable liver damage, when widespread immune activation would overwhelm the ability to detect any initial triggers of inflammation.

We used this predictable time window to investigate for the earliest events of immune activation. We recruited patients stopping NA therapy and collected blood and liver fine-needle aspirates (FNAs) at the time of stopping NA and 4 weeks post-NA withdrawal, when HBV replication rebounded but ALT remained normal. Our data demonstrate that the earliest signatures of inflammation were driven by type I IFN, were only detectable within the liver and that inflammation could be further enhanced by the expression of macrophage migration inhibitory factor (MIF).

## RESULTS

### Study design and patient cohort

11 HBeAg-CHB patients were enrolled under an ancillary study to the Hepatitis B Research Network immune active NA-withdrawal cohort. Patients were on tenofovir disoproxil fumarate (TDF) for 192 weeks or 24 weeks of pegylated-IFN-α then 168 weeks of TDF, prior to a protocol defined stop in treatment. All patients were HBeAg-negative, anti-HBe-positive and had undetectable HBV DNA and normal ALT levels without cirrhosis at the time of treatment withdrawal (Supplementary table 1). Blood was collected from all 11 patients while liver FNA samples were collected from 6 patients at the time of NA withdrawal (Week 0) and 4 weeks post-withdrawal (Week 4) (Fig. 1A). However, one patient was excluded due to elevated ALT levels at baseline, before NA-withdrawal. Upon stopping therapy, all patients experienced viral rebound and displayed moderate to severe ALT elevation (101 – 1914 U/L; Fig. 1B). A hepatic flare was defined as ALT >5x upper limit of normal (ULN; 35 IU/mL for males, 25 IU/mL for females). Immunological assays performed on each patient were summarized (Supplementary table 2).

**Figure 1:**
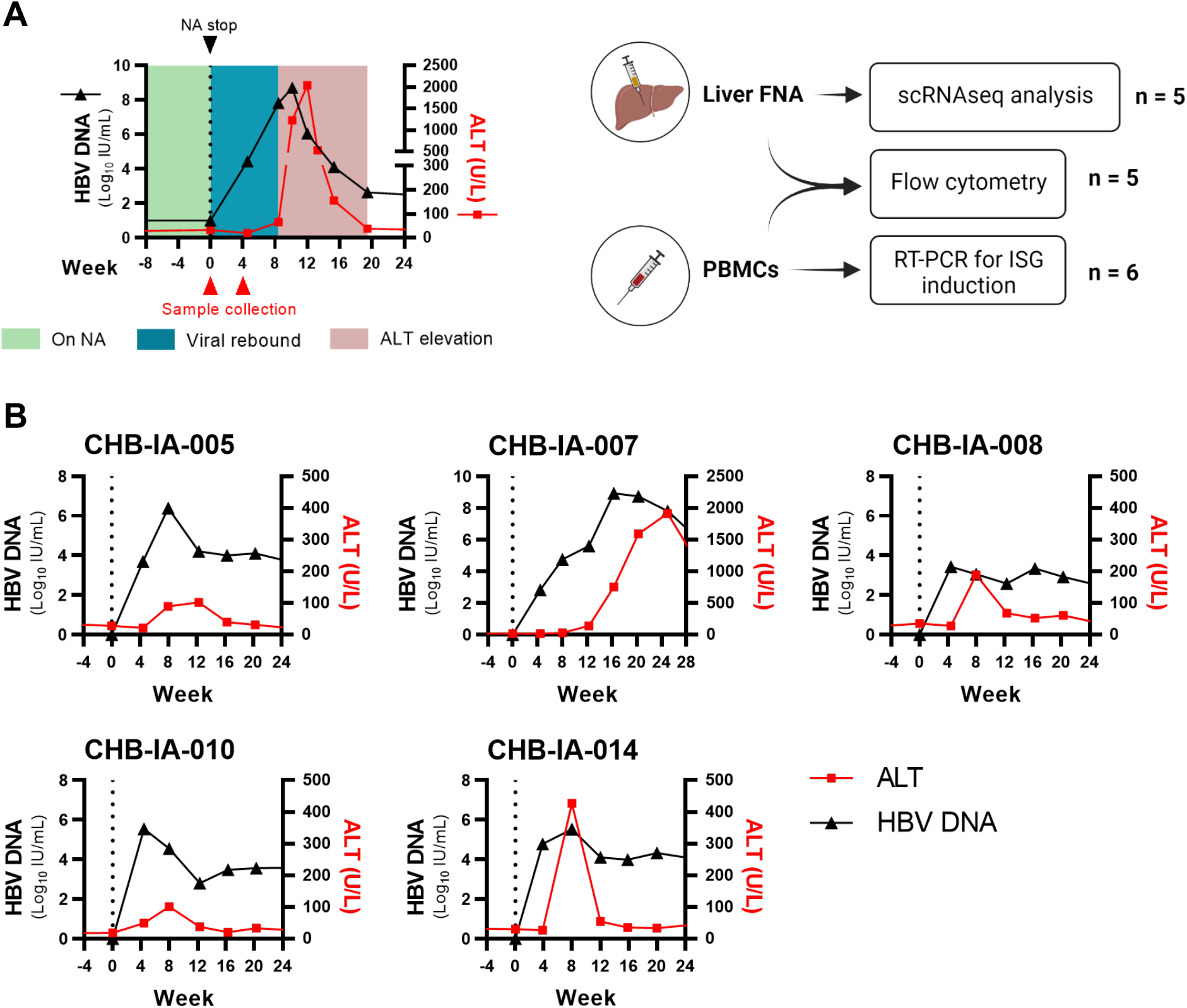
Study design and patient clinical data following 192 weeks of NA therapy. (A) Study design includes 11 CHB patients withdrawing from NA-therapy. Liver FNAs were collected from 5 patients at baseline and viral rebound (week 4 post-withdrawal). (B) CHB patients were on 192 weeks of NA therapy before protocol defined stop. HBV DNA (black) and serum ALT levels (red) were measured for each patient for 28-weeks following NA-withdrawal (dotted vertical line).

### Characterization of longitudinal Liver FNA samples after nucleoside analogue withdrawal

To characterize the liver FNAs, single-cell RNA sequencing (scRNA-seq) was performed on longitudinal FNAs (n = 5). UMAP plots were generated and labelled by patient (Fig. 2A) and timepoint (Fig. 2B) to demonstrate the distribution of captured events. A total of 18,468 cells (following quality control procedures) were captured and clustered into 19 discrete cell populations (Fig. 2C). Cluster identification was performed with dot plots using known cell type markers (Supp. Fig 1A). Clusters 0, 1, 6, 9 and 13 were classified as NK cell clusters for their expression of *KLRB1* and *KLRF1*. Clusters 2, 3, 4, 5 and 7 were T cells evidenced by robust *CD3D* expression. Cluster 10 was the sole B cell cluster detected. Clusters 8, 11, 12, 15 and 17 were broadly classified as myeloid cells for their high expression of HLA transcripts. Cluster 18 were hepatocytes, cluster 14 were proliferating cells, and cluster 16 were liver sinusoidal endothelial cells (LSECs). Feature plots highlight marker expression patterns within the UMAP to visually assist with cluster identification (Supp. Fig 1B). Further cellular markers were consolidated to derive specific cluster identities (Fig. 2D). Cell frequencies obtained from scRNA-seq data were compared between baseline and week 4 (Fig. 2E). No significant differences in cell frequencies were found in any cluster between time points. This was validated via flow cytometry data between baseline and week 4, which did not find any significant differences in frequencies of T cells (CD4+, CD8+, γδ, and MAIT cells), B cells, NK cells, monocytes, and neutrophils in either compartment (Supp. Fig. 2 & 3).

**Figure 2:**
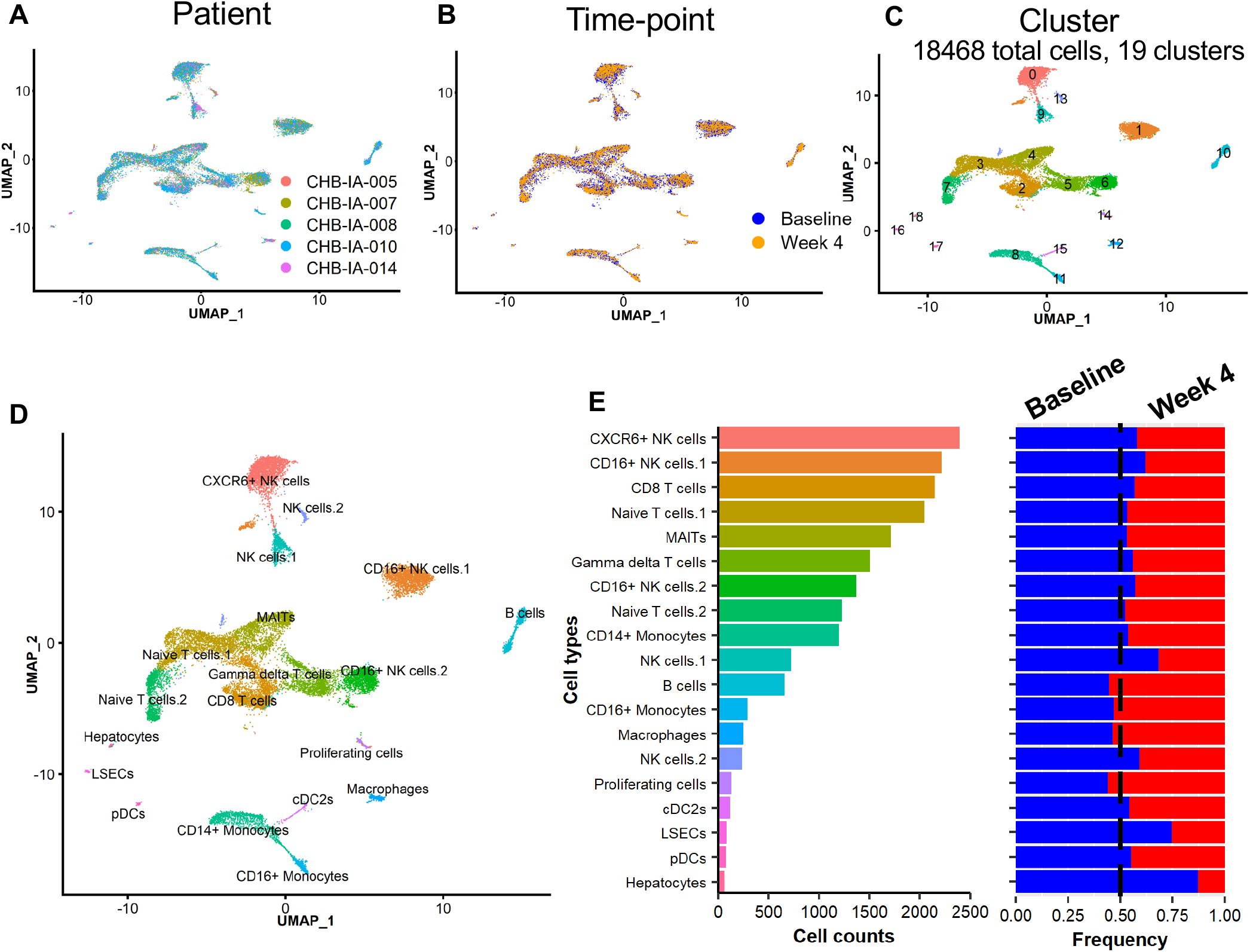
Single-cell RNA-sequencing analysis of intrahepatic immune compartment among CHB patients withdrawing NA-therapy. UMAP plots were generated to distinguish events between (A) patient, (B) timepoint, and (C) cell cluster. (D) UMAP plot annotated with derived cluster identities. (E) Cluster cell numbers obtained in the integrated dataset. Cluster frequencies were calculated for each patient between baseline (blue) and week 4 viral rebound (red).

These data demonstrate that we captured the liver samples prior to the characteristic immune cell infiltrates that are associated with liver damage. Therefore, we anticipate that any transcriptional changes observed in liver immune cells will represent the earliest events in the inflammatory cascade.

### Type I interferon signatures and MIF upregulated at viral rebound following NA-withdrawal

For our initial analysis, we pursued a top-down approach by compiling cell clusters into major cell subsets and measuring for broad gene changes to first identify clusters or genes of interest pertaining to immune activation (Fig. 3A). Clusters 14 (proliferating cells), 16 (LSECs), and 18 (Hepatocytes) were excluded given their sparse and inconsistent proportions captured across different patients and timepoints. Differential gene expression analysis was performed using R package *Seurat*, and differentially expressed genes (DEGs) were quantified for each major immune subset between baseline and 4 weeks post NA-withdrawal, the time of viral rebound. Myeloid cells displayed the most DEGs 4 weeks after stopping therapy with 33 DEGs. T, NK, and B cells had 25, 20 and 15 DEGs respectively (Fig. 3B; Supplementary table 1). DEGs were visualized for each major cell subset through volcano plots (Fig. 3C). Of note, *IFITM1* was observed to be significantly upregulated in NK, T and myeloid cells at week 4 viral rebound. This was accompanied by the upregulation of other interferon stimulated genes (ISGs), including *IFITM3, ISG20, ISG15*, and *STAT1*. Macrophage migration inhibitory factor (*MIF*) and its cell surface receptor, CD74, were also observed to be significantly upregulated upon viral rebound among NK and T cell subsets (Fig. 3C). We tested whether intrahepatic ISG induction was reflected in the peripheral compartment. RT-PCR for *ISG15, MX1*, and *OAS1* transcripts in total peripheral blood mononuclear cells (PBMCs) did not reveal significant differences between baseline and viral rebound (Supp. Fig. 4). This furthers the notion that the immunological impact of HBV viral rebound is restricted to the liver compartment.

**Figure 3:**
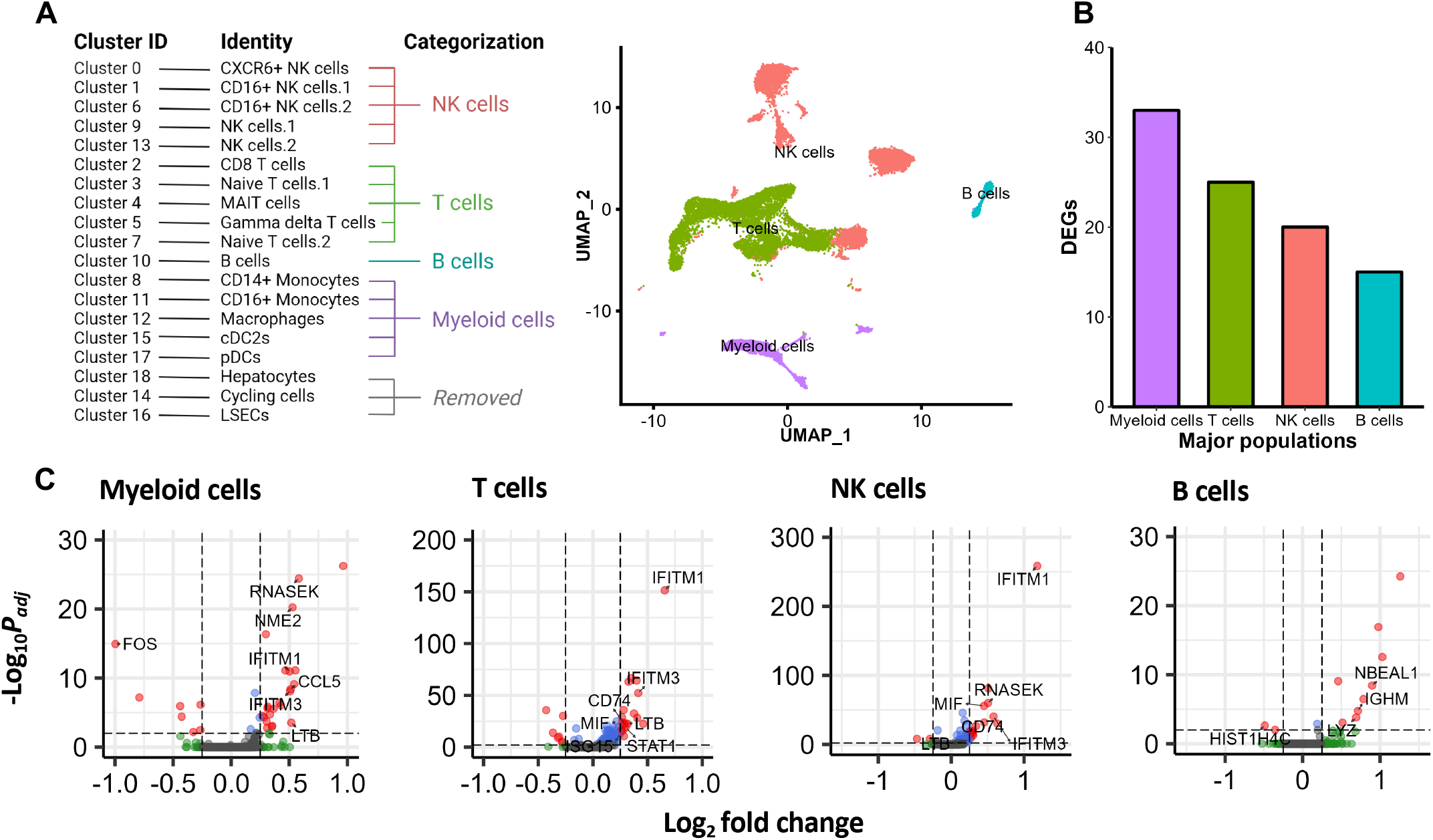
Differentially expressed genes upon viral rebound identified in major intrahepatic immune cell populations. (A) Major immune cell subsets were clustered together as outlined. Non-immune cells (clusters 14, 16, 18) were excluded from DE analysis. (B) DEGs between baseline and week 4 viral rebound were tabulated for each immune cell subset. (C) Volcano plots visualize DEGs (red) for each major immune subset. DEGs are defined using thresholds of >0.25 log_2_ fold-change (vertical lines) and adjusted p-value <0.05 (horizontal lines).

**Figure 4:**
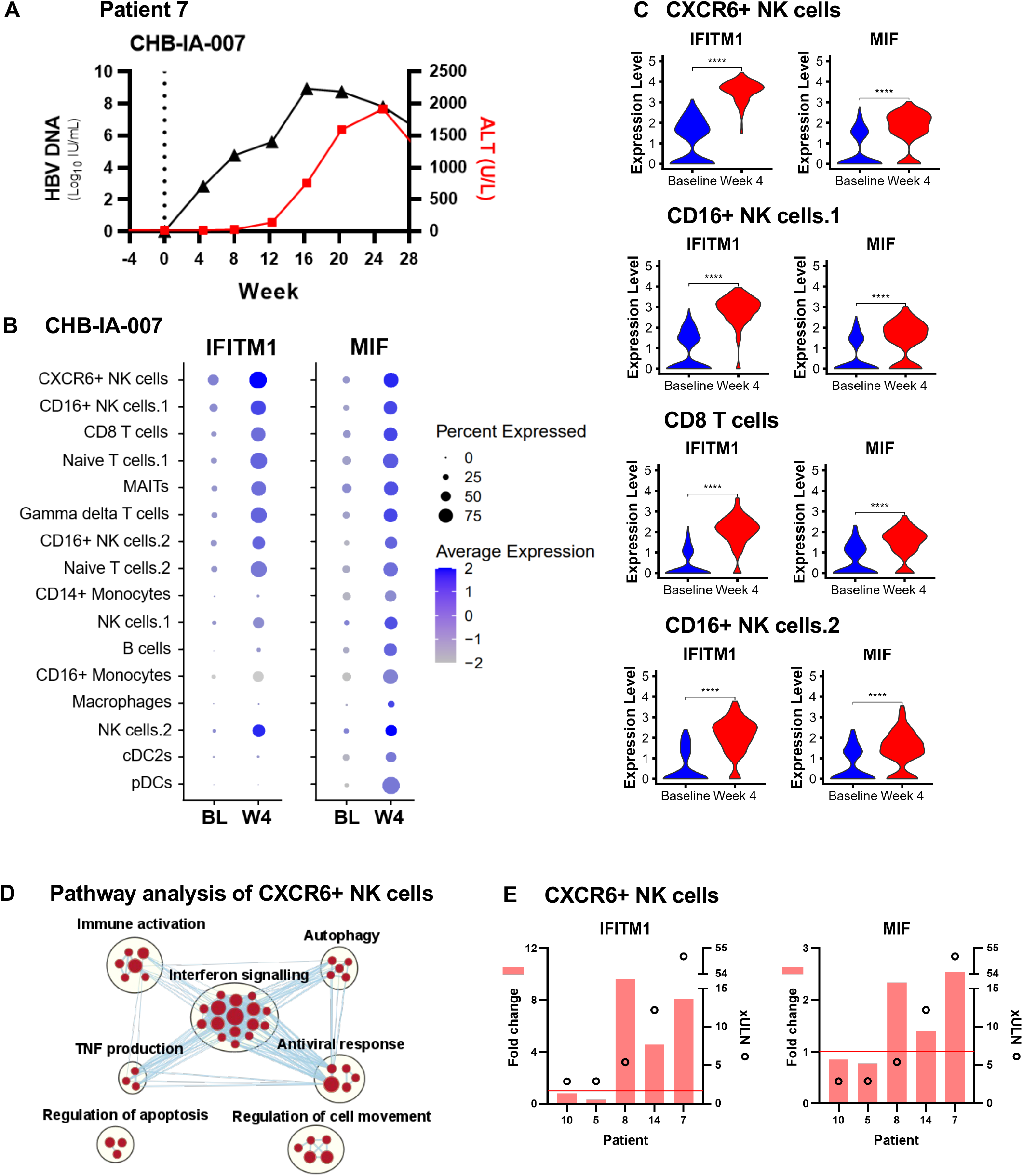
Type I interferon signatures and MIF upregulated at viral rebound following NUC withdrawal. (A) Clinical parameters of patient 7, who experienced the highest peak ALT flare of 54.68xULN at week 24 post-withdrawal. (B) Dot plots of average *IFITM1* and *MIF* expression across patient 7’s cell clusters between baseline and viral rebound. (C) Violin plots of average *IFITM1* and *MIF* expression at baseline (blue) and week 4 (red). (D) GSEA of CXCR6+ NK cells were performed for enriched pathways between baseline and week 4. Enrichment maps were visualized using Cytoscape (v3.8.2). No significantly downregulated pathways were found. (E) Fold change of *IFITM1* and *MIF* expression (red bars) among CXCR6+ NK cells against each respective patient’s peak ALT (empty circles). Wilcoxon tests were conducted to compare *IFITM1* and *MIF* expression levels between baseline and week 4 (**** *P* < .0001).

Overall, we observed modest changes in gene expression, confirming that we captured the earliest events in the inflammatory cascade. The significant DEGs that we observed, across the broad cell clusters, point to a type I IFN response and the up-regulation of MIF, a cytokine previously demonstrated to enhance inflammation in pathogenic environments, which may serve as one of the earliest markers of liver damage.

### ALT flares were preceded by upregulation of IFITM1 and MIF at viral rebound

Having identified a broad type I IFN signature across immune cell types (Fig. 3C), we refined our analysis to assess changes in gene expression in individual cell types defined in figure 1F. To achieve the highest sensitivity for this refined analysis, we initially analyzed the longitudinal samples from patient CHB-IA-007, who experienced the most severe ALT flare at 54.68xULN at 24 weeks post-withdrawal (Fig. 4A). Given *IFITM1* and *MIF* were among the most significant DEGs in figure 2, we measured changes in their expression between baseline and 4 weeks post NA-withdrawal in each cell cluster (Fig. 4B). *IFITM1* displayed significant up-regulation in lymphocyte populations while *MIF* was up-regulated in all immune cell clusters. *IFITM1* was not highly expressed in myeloid populations (Fig. 4B) but other IFN stimulated genes (ISGs), such as IFITM3, were significantly elevated in myeloid cells (Fig. 3C). Violin plots highlighted clusters of CXCR6+ NK cells, CD16+ NK cells.1, CD8 T cells, and CD16+ NK cells.2 as significantly up-regulating *IFITM1* and *MIF* at viral rebound (Fig. 4C, P < .0001). Analyses for other patients were also included (Supp. Fig 5).

**Figure 5:**
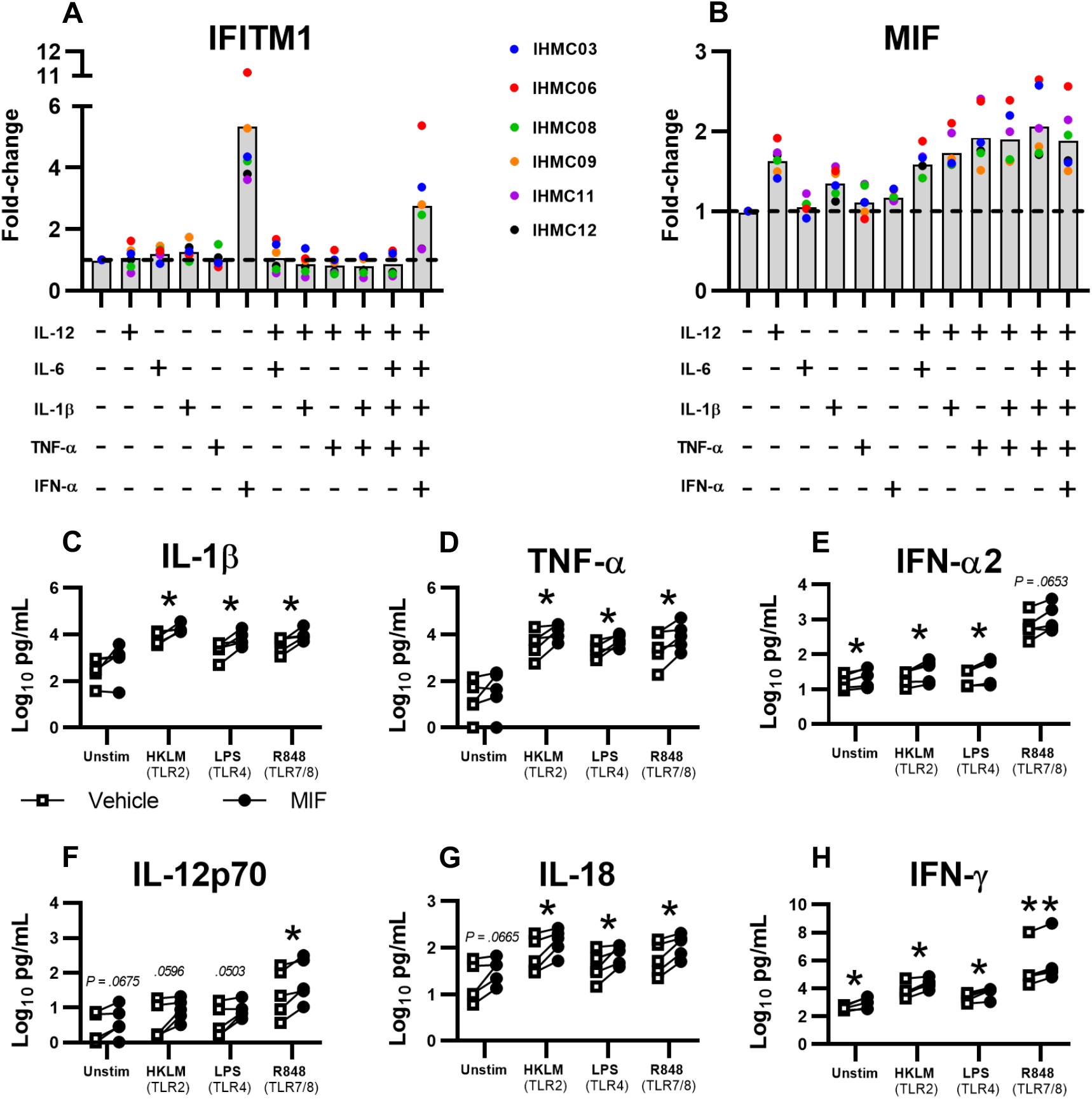
In vitro induction of *IFITM1* and *MIF* and subsequent impact of recombinant MIF protein on cytokine production following TLR stimulation. (A-B) Healthy donor IHMCs (n = 6) were stimulated for 24hr in various cytokine conditions and RT-PCR was performed to quantify induction of *IFITM1* and *MIF* transcripts. Dotted lines highlight fold-change of 1. (C-H) IHMCs (n = 5) pre-stimulated for 6hr in recombinant MIF (filled circles) or vehicle control (empty squares) were stimulated (or unstimulated) for 18hr with TLR agonists, HKLM (TLR2 agonist), LPS (TLR4 agonist) and R848 (TLR7/8 agonist). Cell culture supernatants were collected and quantified for cytokine levels using LEGENDPlex™ Human Inflammation Panel 1 kit (* *P <* .05; ** *P <* .005).

Because of the significant upregulation of both sentinel genes in CXCR6+ NK cells, we used this cluster to perform pathway analysis and determine immune processes enriched at viral rebound (Fig. 4D, Supplementary table 2). At viral rebound, there was a significant enrichment pertaining to major gene sets of immune activation, including tumor-necrosis factor (TNF) production, interferon signaling, antiviral responses and autophagy. The top 3 enriched pathways in GSEA analysis identified robust type I interferon signaling at viral rebound compared to baseline NA-withdrawal (not shown). Regulation of apoptosis and reduced cell movement were primarily attributed to *MIF* (no contributions from ISGs), highlighting the various pleiotropic functions of MIF. There were no downregulated pathways at viral rebound that achieved significance in the CXCR6+ NK cell cluster.

Having defined that the enriched pathways primarily derived from robust *IFITM1* and *MIF* expression, we plotted *IFITM1* and *MIF* fold-change across all 5 patients within the cohort at viral rebound with their peak ALT observed 8 – 24 weeks post stopping NA therapy. Among patients experiencing flares (≥5xULN, patients 8, 14 and 7), *IFITM1* and *MIF* were upregulated at viral rebound prior to the detection of elevated ALT in the serum (Fig. 4E). Patients with moderate ALT elevation (<3xULN; patients 10 and 5) demonstrated no change in *IFITM1* and *MIF* at viral rebound. Comparisons among CD16+ NK cells.1, CD8 T cells, and CD16+ NK cells.2 clusters were performed as well and were consistent with CXCR6+ NK cells (Supp. Fig. 6). These data suggest that an early type I IFN signature and MIF up-regulation are evident across multiple immune populations in the liver of CHB patients prior to elevation of ALT ≥5xULN. Serum from the Toronto NA STOP study^15,16^ were measured by ELLA ProteinSimple assays for MIF levels and IFN-α to try to validate intrahepatic transcriptional changes. However, IFN-α was undetectable in the serum and MIF levels did not distinguish between patients with 0-2xULN, 2-5xULN and ≥5xULN ALT flares at viral rebound (Supp. Fig. 7). These data further support the intrahepatic transcriptional changes observed in bulk immune cell populations and provide further evidence that serum cytokine detection is not sensitive enough to predict outcome after stopping NA therapy^15^.

### Regulation of IFITM1 and MIF expression

The observation that *IFITM1* and *MIF* were broadly upregulated across most clusters suggests that intrahepatic cytokines are likely inducing the expression of these genes. Yet peripheral serum cytokine data from NA-withdrawing patients did not reveal differences between baseline and viral rebound^8^ (Supp. Fig. 7). *IFITM1* is an ISG known to be induced by type I IFN, whereas MIF is reported to be induced by glucocorticoids^17^, bacterial endotoxins and inflammatory cytokines, including TNF-α and IFN-γ^18,19^. Initially, T cells were thought to be the main source of MIF^20–22^, but monocytes, macrophages, dendritic cells and B cells have since been demonstrated to express MIF^20,23–27^. MIF is also expressed in tissues in direct contact with the natural environment, including the lung, the epithelial lining of the skin and the gastrointestinal tract^27–31^.

To validate the role of cytokines in *IFITM1* and *MIF* induction, we stimulated healthy donor intrahepatic mononuclear cells (IHMCs) in vitro with a panel of inflammatory cytokines speculated to stimulate *IFITM1* and *MIF*. IHMCs (n = 6) were stimulated for 24hrs with IL-12p70, IL-6, IL-1β, TNF-α and IFN-α and assessed gene upregulation via RT-PCR (Fig. 5A, B). Upregulation of *IFITM1* was induced solely by IFN-α (Fig. 5A). IL-12p70 alone induced the strongest upregulation of *MIF* (mean FC = 1.65x) compared to other single cytokine conditions, but was further enhanced in combination with additional cytokines, particularly TNF-α. The greatest *MIF* induction was observed with a combination of IL-12p70, IL-6, IL-1β and TNF-α (Fig. 5B; 2.08x mean FC). Our findings suggest that, at a minimum, type I IFN and IL-12p70 are present in the liver to induce *IFITM1* and *MIF*. We tried to measure *IL-12, IFN-α, IFN-β*, and *IFN-λ1/2* transcripts in patient FNA samples but these cytokines were not detectable at either timepoint via scRNA-seq nor via RT-PCR (not shown).

### MIF enhances cytokine production upon TLR stimulation

Elevated MIF protein levels have been associated with more severe cases of sepsis, rheumatoid arthritis, and systemic lupus erythematosus^32–35^. MIF stimulates the release of proinflammatory cytokines such as TNF-α, IL-1, IL-6, IL-8, and IL-12 from macrophages^33,36,37^. Therefore, published data suggest that MIF could enhance the inflammatory response, driving liver damage in patients stopping NA therapy. To determine if the presence of MIF leads to greater induction of cytokine production, we stimulated IHMCs with Toll-like receptor (TLR) agonists in the presence of recombinant MIF. IHMCs (n = 5) were incubated for 6hr with recombinant MIF, followed by stimulation with TLR agonists HKLM (TLR2 agonist), LPS (TLR4 agonist) and R848 (TLR7/8 agonist) for 18hr. IL-1β secretion significantly increased during TLR2 (P = .0294, mean fold change (FC) = 2.42x), TLR4 (P = .0245, mean FC = 3.11x) and TLR7/8 stimulations (P = .0324, mean FC = 2.54x) (Fig. 5C). TNF-α secretions followed a similar pattern (P = .0442, mean FC = 2.26x; P = .0324, mean FC = 2.42x; P = .0396, mean FC = 2.78x respectively) (Fig. 5D). Neither analyte was induced by MIF alone. Interestingly, IFN-α2 secretion was induced by MIF alone (P = .0187, mean FC = 1.41x), and significantly enhanced after stimulation with HKLM (P = .0334, mean FC = 1.73x) and LPS (P = .0340, mean FC = 1.82x), but not with R848 (P = .0653, mean FC = 1.76x) (Fig. 5E). MIF significantly enhanced IL-12p70 secretion after R848 stimulation (P = .0184, mean FC = 2.05x) (Fig. 5F). TLR2/4/7/8 stimulation with MIF improved IL-18 secretion (P = .0210, mean FC = 1.74x; P = .0376, mean FC = 1.65x; P = .0107, mean FC = 1.71x) and IFN-γ secretion (P = .0131, mean FC = 2.08x; P = .0153, mean FC = 2.77x; P = .0015, mean FC = 4.08x) (Fig. 5G-H). These data show that MIF can induce type I and type II IFNs in the absence of inflammatory signals and enhance cytokine production following activation through TLR stimulation.

## DISCUSSION

Spontaneous ALT flares in CHB patients are unpredictable, making it virtually impossible to capture the earliest events in the pathogenic process. NA-withdrawal provides a predictable window of time to study early inflammatory events that subsequently lead to liver damage. ScRNA-seq of CHB patient liver FNA samples provided a high-resolution, single-cell perspective of early intrahepatic immune processes activated at viral rebound following NA-withdrawal. At this early time point, we did not observe changes in cell frequencies, indicating that the inflammatory cellular infiltrate associated with liver damage had not yet occurred. Instead, we found upregulation of numerous ISGs such as *IFITM1, IFITM3, ISG20, ISG15*, and *STAT1*, in addition to pro-inflammatory cytokine *MIF. IFITM1* and *MIF* induction occurred weeks before the onset of clinically detectable liver damage and were greatest among NA-withdrawing patients who experienced ALT flares ≥5xULN. Broad upregulation of these marker genes suggested that immune activation was mediated by cytokines, which we validated using IHMCs from healthy donors. We further demonstrated the inflammatory potential of MIF through its ability to induce type I and II IFNs and other inflammatory cytokines that can further perpetuate the inflammatory response among TLR-stimulated (and unstimulated) intrahepatic immune cells.

The activation of type I IFN elicits immune responses against viral infections and has been observed during liver damage in chronic HBV infection^7–9,38–40^. Acute HBV-infected chimpanzee studies correlated the induction of various ISGs to peak ALT, highlighting the contribution of ISGs to the inflammatory response during hepatic flares^39^. IFN-α is also known to enhance immune recognition through upregulating MHC expression^41^. Our study reports the upregulation of several HLA genes alongside ISG induction during viral rebound, providing further evidence for the activation of type I IFN responses at this timepoint. Furthermore, IFN-α was shown to drive CD8 T cell activation and propagate liver damage in a mouse model of non-alcoholic fatty liver disease^42^. These data support our observation that upregulation/appearance of type I IFNs is the earliest event in the inflammatory cascade leading to liver damage.

We were not able to identify the source of type I IFN, or IL-12, from our dataset. Cytokines were poorly captured in the scRNA-seq data and parenchymal cells were not highly abundant because of quality control metrics used to filter the data, which primarily focused on immune cells. Hepatocytes have the potential to serve as the source of type I IFNs through RIG-I, which has been demonstrated as a sensor for pre-genomic RNA in infected hepatocytes^43^. Alternatively, the rapid rebound in viral replication has the potential to induce cellular stress and the production or release of endogenous danger signals, damage-associated molecular patterns (DAMPs). DAMPs could then promote activation of innate immune cells and initiate liver inflammation through pattern recognition receptor (PRR)-mediated cytokine secretion. Further hepatocyte lysis could perpetuate this inflammatory state and manifest as hepatic flares. Thus, this cascade has the potential to account for both type I IFN and IL-12 to be produced by either liver macrophages or sinusoidal monocytes and dendritic cells to broadly induce both ISGs and MIF expression among intrahepatic immune cells. However future studies focused on identifying the cell-types and the PRRs activated during viral rebound are required.

The identification of MIF potentially playing a role in HBV-associated liver inflammation introduces a new pathway of immune activation. MIF protein is an inflammatory cytokine implicated in the pathogenesis of several inflammatory diseases including sepsis, rheumatoid arthritis, and systemic lupus erythematosus^32–35^. Although MIF remains poorly studied in the context of CHB, elevated serum MIF levels have recently been associated with disease severity among patients with decompensated cirrhosis and acute-on-chronic liver failure^44^. MIF is known to be a critical mediator of inflammation^17,45,46^, stimulating the release of TNF-α, IL-1 and IL-6 from macrophages^36^ and IL-1 and IL-8 from synovial fibroblasts^33,37^. The MIF receptor, CD74, is a prominent cell surface activation marker present on lymphoid cells and is observed during peak ALT levels in HBV-infected chimpanzees^39^. Additionally soluble CD74 (sCD74), which acts as a decoy receptor to sequester extracellular MIF and neutralize the effects of MIF signaling, has been shown to correlate with improved clinical outcomes^44,47–49^. Indeed, high levels of MIF coupled with low sCD74 levels were recently reported to be predictive of poorer 90-day transplant-free survival among patients with decompensated cirrhosis and acute-on-chronic liver failure^44^.

In our study, MIF alone could induce cytokine production and further enhanced secretion of various inflammatory cytokines upon TLR stimulation in vitro. We found that *MIF* expression was induced by inflammatory cytokines, particularly by IL-12, which suggests that MIF can potentially function in a positive feedback loop to perpetuate various inflammatory processes and enhance immune activation. However, we have yet to determine if IL-12 is the initial trigger for MIF induction since the IL-12 receptor will be largely restricted to lymphocytes and we observed almost ubiquitous MIF transcript up-regulation across immune cell types. If there is a ubiquitous danger signal that induces MIF, its early expression could set the inflammatory cascade into motion with its ability to stimulate the production of MIF and type I IFNs. The limited reagents available to measure human MIF protein presents practical hurdles but, given the clinical relevance of MIF/sCD74 in inflammation, MIF could represent a promising prognostic marker for severe ALT flares following NA-withdrawal or serve as a potential target to modulate liver damage among CHB patients.

Our study investigated transcriptional activation of immune cells at viral rebound. The primary goal was to identify peripheral biomarkers that could predict the severity of subsequent liver damage, however we were unable to translate intrahepatic signatures to peripherally detectable markers. Cell frequencies did not change between timepoints and ISG induction was only detectable in the liver. Intrahepatic analyses identified type I IFN and MIF as the earliest up-regulated cytokines, particularly in patients that experienced ALT flares ≥5xULN. Induction of these genes was achieved in vitro using a panel of cytokines associated with innate activation, suggesting that either HBV-derived pathogen-associated molecular patterns (PAMPs) or endogenous DAMPs represent the earliest “danger” signal. The ability of MIF to enhance cytokine production provides positive feedback mechanisms to propagate liver damage. Understanding these early immune events driving hepatocyte killing is critical to understand the outcome of liver damage and to better inform clinical decisions for the optimal management of CHB flares.

## MATERIALS AND METHODS

### Ethical statement and human subjects

The Hepatitis B Research Network (HBRN) Cohort Studies were approved by the research ethics boards or institutional review boards of all participating sites. Samples used for the current study were collected prospectively at the Toronto Centre for Liver Disease following review and approval by the HBRN ancillary study steering committee. All patients provided written informed consent. Patient clinical data is as follows (Supplementary table 3) as well as a summary of assays performed (Supplementary table 4)

Our validation cohort consisted of sera collected from NA-withdrawing patients in the Toronto STOP study (ClinicalTrials.gov, Identifier: NCT01911156). Samples were collected at the Toronto Centre for Liver Disease (University Health Network, Canada) from May 2016 to May 2018. This study was approved by the research ethics board of University Health Network in Toronto and performed in concordance with Good Clinical Practice guidelines and the Declaration of Helsinki (2013). All patients provided written consent.

### Flow cytometry

Fluorophore-conjugated antibodies were used to discern immune cell populations in peripheral whole blood and liver FNA samples with the following flow schemes (Supp. Fig. 2, 3). Dead cells were stained with eFluor 520 (eBioscience) in PBS for 10 min. at room temperature. For surface marker staining, cells were incubated with antibodies in 4^°^C for 30 min. (Supplementary table 5). Cells stored in 1% paraformaldehyde in preparation for FACS analysis.

### Liver FNA processing

Liver FNA were performed by a hepatologist. A suitable location was determined using ultrasound and anesthetized with lidocaine. 25-gauge needles were used for puncture and cells were aspirated into a 10mL syringe containing 0.5mL of RPMI medium (Gibco). After collection, the needle was flushed with an additional 0.5mL RPMI to collect remaining cells. A total of four liver FNA passes were taken from each patient at each timepoint. The exact volume of each FNA pass was documented and the optical density (OD) of each pass was measured to assess for peripheral blood content. For scRNA-seq, red blood cells (RBC) were removed by 5 min. incubation with RBC Lysis Buffer (BioLegend) and subsequent 10x dilution with PBS. Cells were washed twice, counted, and 20,000 cells per sample resuspended in 50µL PBS + 0.04% BSA for scRNA-seq. Any remaining sample was cryopreserved in Knockout Serum Replacement (Gibco) + 10% DMSO.

### 10x sample preparation and cDNA library preparation

Samples were prepared as outlined by 10x Genomics Single Cell 3’ Reagent Kits v2 user guide. Following counting, the appropriate volume for each sample was calculated for a target capture of 2000 or 3000 cells. Samples that were too low in cell concentration as defined by the user guide were washed, re-suspended in a reduced volume prior to loading onto the 10x single cell A chip. After droplet generation, samples were transferred onto a pre-chilled 96-well plate (Eppendorf), heat sealed and incubated overnight in a Veriti 96-well thermos cycler (Thermo Fisher). The next day, sample cDNA was recovered using Recovery Agent provided by 10x and subsequently cleaned up using a Silane DynaBead (Thermo Fisher) mix as outlined by the user guide. Purified cDNA was amplified for 12 cycles before being cleaned up using SPRIselect beads (Beckman). Samples were diluted 4:1 (elution buffer (Qiagen): cDNA) and run on a Bioanalyzer (Agilent Technologies) to determine cDNA concentration. cDNA libraries were prepared as outlined by the Single Cell 3’ Reagent Kits v2 user guide with modifications to the PCR cycles based on the calculated cDNA concentration.

### Sequencing

The molarity of each library was calculated based on library size as measured bioanalyzer (Agilent Technologies) and RT-PCR amplification data (Sigma). Samples were pooled and normalized to 10nM, then diluted to 2nM using elution buffer (Qiagen) with 0.1% Tween20 (Sigma). Each 2nM pool was denatured using 0.1N NaOH at equal volumes for 5 min. at room temperature. Library pools were further diluted to 20pM using HT-1 (Illumina) before being diluted to a final loading concentration of 14pM. 150µL from the 14pM pool was loaded into each well of an 8-well strip tube and loaded onto a cBot (Illumina) for cluster generation. Samples were sequenced on a HiSeq 2500 (Illumina).

### Preprocessing of scRNA-seq data

Raw sequencing data (bcl files) were converted to fastq files with Illumina bcl2fastq, version 2.19.1 and aligned to the human genome reference sequence [http://cf.10xgenomics.com/supp/cell-exp/refdata-cellranger-GRCh38-1.2.0.tar.gz]. The CellRanger (10x Genomics) analysis pipeline was used to generate a digital gene expression matrix from this data. Resulting gene-barcode matrices were subjected to further quality control filtering, data preprocessing, scaling and normalization by *Seurat* package (v4.0.0). Cells with a very small library size (<200) or high mitochondrial DNA content (>10%) were removed. Genes present in less than 3 cells were filtered out. After applying these quality control criteria, 18,468 single cells remained for downstream analyses.

### Cell clustering, differential expression, and gene set enrichment analyses

The *Seurat* (v4.0.0) package was used for the analysis of the data sets. Clustering was performed using the Louvain algorithm with resolution set to 0.3. To visualize these data, we applied a uniform manifold approximation and projection (UMAP) algorithm to perform nonlinear dimensionality reduction (Fig. 2B-C). Cluster identities were determined manually using a compiled panel of known surface markers for each intrahepatic cell population (Supp. Fig. 1A-B). Differentially expressed genes (DEG) of major cell subtypes were calculated between baseline (week 0) and viral rebound (week 4) using the FindMarkers function (test.use = “wilcox”) with a log_2_ fold-change threshold of >0.25, and adjusted p-value <0.05 (Supplementary table 1). Volcano plots were visualized using the *EnhancedVolcano* package (Fig. 3C).

Gene rank lists were compiled in *Seurat* (Supplementary table 2). Cellular pathways enriched were further investigated using the Gene Set Enrichment Analysis (GSEA) software from the Broad Institute (v4.0.3).

Human_GOBP_AllPathways_no_GO_iea_August_01_2021_symbol.gmt from [http://baderlab.org/GeneSets] was used to identify enriched cellular pathways in GSEA analysis and enrichment maps were visualized using Cytoscape (v3.8.2). An adjusted p-value <0.05 and q-value <0.1 were used as significance thresholds for nodes. Edge cutoffs were set at <0.3 and highly related pathways were grouped into a theme and labeled by AutoAnnotate (v1.2).

### IHMC processing

IHMCs were isolated from living donor liver transplantation perfusions at Toronto General Hospital. Prior to transplantation, livers were perfused with 1 L of cold University of Wisconsin (UW) solution (ViaSpan). Perfusates were collected and centrifuged for concentration in HBSS + 2 U/ml heparin, to a final volume of 50-100mL and subjected to density centrifugation using Lymphoprep solution (StemCell). Cells were counted and cryopreserved in Knockout Serum Replacement + 10% DMSO.

### In vitro stimulation of donor IHMCs

Cryopreserved IHMCs were thawed and resuspended at 2 × 10^6^ IHMCs/mL in Aim-V media. 1 × 10^6^ IHMCs were plated in each 48-well and incubated with either IL-12p70 (100ng/mL), IL-6 (100ng/mL), IL-1β (100ng/mL), TNF-α (100ng/mL), or IFN-α (100U/mL) for 24h at 37^°^C in 5% CO_2_. All recombinant human cytokines were purchased from GoldBio (Massachusetts, USA). Following stimulation, cell pellets were collected by centrifugation at 300xg for 5min. and resuspended in 300µL RLT buffer (Qiagen) + 1% β-mercaptoethanol.

For multiplex cytokine analysis, 5 × 10^5^ IHMCs were plated in each 96-well and incubated in 75µL complete R10 media with rhMIF (R&D Systems; 1mg/mL) or vehicle (PBS + 0.2% BSA) for 6h at 37^°^C in 5% CO_2_. Complete R10 media contains: 50x MEM Amino Acids without L-glutamine (Corning), 100x MEM Nonessential Amino Acids (Corning), 100mM Sodium Pyruvate solution (Corning, working concentration: 1mM), 1M HEPES Buffer (Corning, working concentration: 20mM), 100x GlutaMAX solution (Gibco), heat-inactivated FBS (Gibco), 10,000 U/mL Penicillin-Streptomycin (Gibco, working concentration: 100U/mL) and RPMI Medium 1640 without L-glutamine (Gibco). Following MIF exposure, TLR stimulation was performed using HKLM (InvivoGen; 10^5^U/mL), LPS (InvivoGen; 10ng/mL) or R848 (InvivoGen; 100ng/mL) for 18h. Supernatant was analyzed using the LEGENDPlex™ Assay (BioLegend; Human Inflammation Panel 1 (13-plex)) as per manufacturer’s protocols. Results were acquired on a LSR Fortessa (Becton Dickinson) cytometer and analyzed using LEGENDplex™ Data Analysis Software (BioLegend).

### RNA isolation and RT-PCR analysis

RNA extractions were performed as per manufacturer’s protocol (Qiagen). cDNA synthesis was performed using at least 125ng of total RNA and mixing 10x RT Buffer (Thermo), 10x RT Random Primers (Thermo), dNTP Mix 100mM (Thermo), Oligo (dT) 12-18 Primers (Invitrogen), Rnase Inhibitor (Thermo), Rnase-free H_2_O and Multiscribe Reverse Transcriptase (Thermo) to a final reaction volume of 20μL. cDNA synthesis was carried out as follows: 10 min. hold at 25^°^C, 2h hold at 37^°^C for reverse transcription, and a final 5 min. hold at 85^°^C for enzyme inactivation before a perpetual hold at 4^°^C. Quantitative RT-PCR reactions were performed using 2x TaqMan™ Fast Advanced Master Mix (Thermo) with 20x TaqMan™ gene specific assays (Thermo), Rnase-free H_2_O, and 2.5μL of cDNA (at least 12.5ng RNA) to a final reaction volume of 10μL. Amplification was performed as follows: 2 min. hold at 50^°^C, 2 min. at 95^°^C, followed by 40 cycles of 3 seconds at 95^°^C and 30 seconds at 60^°^C. All experiments were performed in triplicates in a QuantStudio 6 (Thermo). Gene expressions were normalized against GAPDH. TaqMan™ Assay probes used are as follows: IFITM1, Hs00705137_s1; MIF, Hs00236988_g1; and GAPDH, Hs99999905_m1; IL-12A, Hs01073447_m1; IFNA2, Hs02621172_s1; IFNB1, Hs02621180_s1 (Thermo); IFNL1, Hs0601677_g1; IFNL2, Hs04193050_gH.

### Serum cytokine analyses

Patient sera were thawed and used as per manufacturer’s protocols for IFN-α ProQuantum (Invitrogen), LEGENDMAX MIF ELISA (BioLegend) and ELLA ProteinSimple kits (Bio-Techne).

## ACKNOWLEDGEMENTS

The authors acknowledge the use of HBRN data as the sole contribution of the HBRN. Additional support was provided to the HBRN by Gilead Sciences, Inc. and Roche Molecular Systems through Cooperative Research and Development Agreements (CRADAs) with the NIDDK, and Roche/Genentech through a Clinical Trials Agreement (CTA) with the NIDDK.

## SUPPLEMENTARY FIGURE LEGENDS

**Supplementary figure 1: Curated list of cell surface markers to identify scRNA-seq clusters**. (A) Dot plots of marker expression levels were used to identify cluster identities. (B) Feature plots visualize a marker’s expression pattern within the UMAP.

**Supplementary figure 2: Flow cytometry gating scheme for peripheral and intrahepatic immune populations**. (A) Peripheral blood and (B) intrahepatic immune populations were assessed via flow cytometry at the point of NA-withdrawal and 4-weeks post-withdrawal.

**Supplementary figure 3: Peripheral and intrahepatic immune cell frequencies did not differ significantly between baseline and viral rebound**. Matched (A) PBMCs and liver FNAs from patients (n = 5) at both timepoints were stained for various immune cell populations and assessed via flow cytometry. No significant changes in cell frequencies were found between timepoints for both peripheral and intrahepatic compartments.

**Supplementary figure 4: RT-PCR analysis of patient PBMCs for ISG expression between flaring and non-flaring patients**. RNA isolated from patient PBMCs was used for RT-PCR analysis for ISG expression, including *ISG15, MX1* and *OAS1*. ISG expression levels were compared between baseline (empty circles) and week 4 viral rebound (filled circles) among flaring and non-flaring patients. No significant differences were observed.

**Supplementary figure 5: *IFITM1* and *MIF* transcript levels across all clusters for each patient**. Dot plots of average *IFITM1* and *MIF* expression across other HBRN-IA patients, visualized by cluster between baseline and viral rebound.

**Supplementary figure 6: *IFITM1* and *MIF* transcript levels in other immune cell clusters against peak ALT levels**. Fold change of *IFITM1* and *MIF* expression (red bars) among (A) CD16+ NK cells.1, (B) CD8 T cells, and (C) CD16+ NK cells.2 against each respective patient’s peak ALT (empty circles).

**Supplementary figure 7: Peripheral sera analyses for MIF and IFN-α levels between flaring and non-flaring patients**. Patient sera from NA-withdrawing patients taken at baseline (empty circles) and week 4 viral rebound (filled circles) were analyzed for MIF levels using ELLA ProteinSimple kits (** *P <* .005).

